# Cystathionine gamma lyase overexpression enhances neovascularization through NAD-dependent mechanisms

**DOI:** 10.1101/2022.09.06.506715

**Authors:** Kevin Kiesworo, Michael R MacArthur, Peter Kip, Thomas Agius, Diane Macabrey, Martine Lambelet, Lauriane Hamard, C.-Keith Ozaki, James R Mitchell, Sebastian Déglise, Sarah J Mitchell, Florent Allagnat, Alban Longchamp

**Author notes:** Correspondence to, Rue du Bugnon 46, 1011 Lausanne, Switzerland. Contributed equally (Florent Allagnat, Alban Longchamp). **Authorship:** KK, FA, MRM, PK, CKO, SJM, JRM and AL designed the study. AL, MRM, SJM, CKO, KK and FA wrote the paper. KK, ML, DM, MRM, PK, TA, FA, and AL performed the experiments. KK, JMC, SD, PK, MRM, TA, SJM, FA and AL analyzed the data. **Disclosure:** The authors have no competing interests.

## Abstract

**Objective:** Hydrogen sulfide (H_2_S) is a proangiogenic gas produced primarily by the transsulfuration enzyme cystathionine-gamma-lyase (CGL). CGL-dependant H_2_S production is required for neovasculariation in models of peripheral arterial disease. However, the benefits of increasing endogenous CGL and its mechanism of action have yet to be elucidated.

**Methods:** 10 weeks old male whole-body CGL overexpressing mice (CGL^Tg^) and wild type littermates (C57BL/6J) were subjected to the hindlimb ischemia model. Functional recovery was assessed through treadmill exercise endurance testing, while ischemic leg perfusion recovery was measured by laser Doppler perfusion imaging and tissue immunohistochemistry. To examine angiogenic potential, aortic ring sprouting assay and post-natal mouse retinal vasculature development studies were performed. Lastly, comparative metabolomics, NAD^+^/NADH analysis, and quantitative real-time PCR were performed on WT and CGL^Tg^ gastrocnemius muscles.

**Results:** The restoration of blood flow upon femoral ligation occurred more rapidly in CGL^Tg^ mice. CGL^Tg^ mice were able to run further and for longer compared to WT mice. In ischemic gastrocnemius, capillary density was increased in mice overexpressing CGL. Endothelial cell sprouting was increased in aorta isolated from CGL^Tg^ mice, especially when cultured in VEGF-only media. Metabolomics analysis demonstrated an increased presence of niacinamide, a precursor of nicotinamide adenine dinucleotide (NAD^+^/ NADH) in the muscle of CGL^Tg^ mice. Finally, CGL overexpression and NMN supplementation improved endothelial cell migration *in vitro*.

**Conclusions:** Taken together, our results demonstrate that CGL overexpression improves the neovascularization of skeletal muscle upon hindlimb ischemia. These effects are mediated by changes in the NAD pathway, which improves endothelial cell migration.

## INTRODUCTION

Peripheral arterial disease (PAD) currently affects more than 200 million people worldwide, and is anticipated to rise with the ageing population (1). PAD can lead to severe complications such as chronic limb threatening ischemia (CLTI) and amputation, and is associated with high rates of cardiovascular events and death. Clinical management of PAD patients aim to improve functional capacity, and maintain limb viability (2). During peripheral artery disease, occlusion of the arteries can trigger a series of compensatory events, such as arteriogenesis and angiogenesis, to restore perfusion in the ischemic tissue (3, 4). One therapeutic strategy that has been explored in this condition is to enhance the formation of new capillary network, through the administration of growth factors and vasculogenic cells, and subsequent activation, proliferation and migration of endothelial cells. Although some success of angiogenic therapy has been reported in young and healthy animal models of acute artery ligation. However, when translated to patient cohorts, these therapies have yet to demonstrate its therapeutic efficacy (3, 5, 6).

In mammals, H_2_S is a ubiquitous redox modifying gasotransmitter that plays numerous physiological roles across various organ systems, including the cardiovascular system (7, 8). H_2_S is produced through the trans-sulfuration pathway by the concerted effort of 3 enzymes: cystathionine-γ-lyase (CGL), cystathionine-β-synthase (CBS), and 3-mercaptopyruvate sulfurtransferase (3-MST) (9). In the cardiovascular system, CGL is thought to be the principal driving force behind the production of H_2_S (10), which has pro-angiogenic properties critical in the context of PAD. Thus, whole body knock-out of CGL impairs recovery in a murine model of PAD (11, 12), while the administration of a H_2_S pro-drug has been shown to improve neovascularization in a porcine PAD model (13). In humans, endogenous H_2_S bioavailability is attenuated in the setting of CLTI and in patients with diabetes-related vascular inflammation (14). Moreover, our group recently demonstrated that circulating H_2_S levels are lower in patients with atherosclerotic disease, and that patients undergoing surgical revascularization with lower H_2_S production capacity have higher rates of post-operative mortality (15).

H_2_S-associated angiogenesis is thought to be driven by stimulation of the VEGF pathway in endothelial cells (EC), via the activation of the VEGFR2 receptor through sulfhydration (16). H_2_S further increases EC glucose uptake and ATP production, which allows rapid energy generation supporting migration during angiogenesis (17). Interestingly, a few studies suggested an interplay between H_2_S and the redox couple nicotinamide adenine dinucleotide (NAD^+^/NADH). NAD^+^ is an essential co-enzyme for cellular redox reactions, such as glycolysis and fatty acid oxidation, making it central in energy metabolism. Moreover, it also functions as a substrate for non-redox enzymes, such as sirtuins and poly (ADP-ribose) polymerases (18, 19). Previously, the H_2_S donor NaHS has been shown to activate the NAD^+^ dependent histone deacetylase Sirtuin 1 (SIRT1) and augment the pro-angiogenic effects of the NAD^+^ precursor nicotinamide mononucleotide (NMN) in primary EC (20). Similarly, post-conditioning using sodium hydroxysulfide (NaHS) improves recovery after cardiac ischemia-reperfusion injury in a rat model through the activation of SIRT1/PGC1α (21). NMN also improves blood flow and increases endurance and vascular remodelling following ischemic injury in elderly mice (20). Moreover, NAD precursors increase the angiogenic capacity of EC *in vitro* (22). In the muscle, supplementation with a NAD^+^ precursor, nicotinamide riboside (NR), also accelerates regeneration in aged and young mice in a model of cardiotoxin–induced muscle damage (23). However, the intricacies of the interaction between H_2_S and NAD^+^ have yet to be elucidated.

To investigate the impact of H_2_S on neovascularization, we utilised a model of limb ischemia and mice overexpressing CGL (CGL^Tg^). Here, we identified CGL as a potent pro-angiogenic trigger *in vivo*, dependent on NAD^+^.

## MATERIALS AND METHODS

Materials and reagents are described in Supplementary Tables.

### Animals

10 to 12-weeks old male WT and CGL^Tg^ on a C57BL/6J genetic background mice were used for all experiments. CGL^Tg^ mice were described previously (24). All mice were housed at standard housing conditions (22 °C, 12 h light/dark cycle), with *ad libitum* access to water and regular diet (SAFE^®^150 SP-25 vegetal diet, SAFE diets, Augy, France). All animal experimentations conformed to the *National Research Council: Guide for the Care* and *Use of Laboratory Animals (National Research Council (U*.*S*.) (25). All animal care, surgery, and euthanasia procedures were approved by the CHUV and the Cantonal Veterinary Office (SCAV-EXPANIM, authorization number 3258 and 3504).

### Hindlimb ischemia (HLI) model

The HLI model was performed as previously described (17). Briefly, mice were anaesthetized with isoflurane (2.5% under 2.5 L O2) and body temperature maintained on a circulating heated pad. Following a 1 cm groin incision, the neurovascular pedicle was visualized under a microscope (LW Scientific, Z2 Zoom Stereo-scope).The femoral nerve and vein were separated from the femoral artery. Two sutures (7-0 silk) were placed above the bifurcation with the epigastric artery, allowing electrocoagulation of the left common femoral artery, proximal to the bifurcation of superficial and deep femoral artery while sparing the vein and nerve. Buprenorphine (0.1 mg/kg Temgesic, Reckitt Benckiser AG, Switzerland) was provided before surgery, as well as a post-operative analgesic every 12h for 36 hours.

### Laser Doppler perfusion imaging (LDPI)

Laser Doppler perfusion imaging (LDPI) was performed as described previously (17). Briefly, mice were kept under isoflurane anesthesia, and body temperature maintained on a circulating heated pad. Once unconscious, we subjected the mouse hindlimbs to the LDPI (Moor Instruments Ltd) system with a low-intensity (2 mW) laser light beam (wavelength 632.8 nm). Hindlimb blood flow was recorded as a 2D color-coded image, with a scan setting of 2 ms/pixel. Blood flow recovery was monitored at baseline, d0 (immediately post-surgery), d1, d3, d5, d7, d10, and d14. LDPI intensity of the ischemic foot was normalized to the corresponding contralateral foot and expressed as ratio between the ischemic/non-ischemic limb.

### Treadmill Test

At 14 days post-HLI, mice were acclimated to the treadmill (Columbus Instruments 6-lane treadmill) at 8 m/min for 5 min for 5 days prior to exercise testing. The next day, mice were run until exhaustion at 5° incline, 8 m/min for 10 min then 10 m/min for 5 min, with a 2 m/min increase in speed every 5 min until exhaustion. Exhaustion was defined as follows: mice remain on the electric grid for 5 seconds or, mice remain on the electric grid for 25 seconds in total, or mice receive more than 25 shocks in 3 min, or mice receive more than 200 shocks in total.

### H_2_S production assay

Measurements of H_2_S production capacity in tissues was performed as previously described (17, 26). Briefly, 80 μg of protein were incubated in 10 mM L-cysteine and 1 mM pyridoxal 5′-phosphate hydrate (Sigma); or, 20 μl of plasma was incubated in 100 mM L-cysteine and 10 mM pyridoxal 5′-phosphate hydrate. This mixture was sealed under lead acetate paper and incubated until black lead sulfide precipitate was detected, but not saturated. Lead sulfide presence was quantified using Fiji software (ver. 1.53f51; http://fiji.sc/Fiji).

### NAD+/NADH quantification

Levels of NAD^+^/NADH were determined using the NAD/NADH Assay kit (Abcam) according to the manufacturer instructions. 20 mg of pulverized tissue were dissolved in the assay buffer from the kit as suggested by the manufacturer and normalized to protein content by Pierce™ BCA Protein Assay Kit (ThermoFisher). 250 μg of protein was used for each reaction. NAD^+^ and NADH concentrations were expressed in ng/mg protein.

### Metabolomics

Polar metabolite profiling was performed as described previously (27). Briefly, 20 mg of mouse gastrocnemius tissue was homogenized on dry ice in 80% methanol and kept at -80°C overnight. Debris were pelleted and the methanol suspension was dried under nitrogen. The resulting pellet was resuspended in water and metabolites were measured using targeted tandem mass spectrometry (LC-MS/MS) with polarity switching and selected reaction monitoring with an AB/SCIEX 6500 QTRAP mass spectrometer.

### Immunohistochemistry (IHC)

IHC was performed on 10 μm frozen sections of gastrocnemius muscle. After 5 min fixation in PFA 4% and rinsing in PBS, immunostaining was performed as previously described (17, 28). Slides were incubated overnight with VE-cad (CD144; BD Pharmingen™; 1:200) rat anti-mouse primary antibody, followed by AlexaFluor 488 anti-rat secondary antibody, and washed and mounted in DAPI-containing Vectashield fluorescent mounting medium. Sections were then scanned with a Ziess Axioscan microscope. VE-cad positive area of whole muscle was quantified blindly using Fiji software (ver. 1.53f51; http://fiji.sc/Fiji). Quantifications were expressed as a percentage of VE-cad-positive area to the total surface area of the gastrocnemius muscle.

### Immunostaining of whole-retina mounts

Eyes from 5-day old pups were harvested and fixed for 2 hours at 4°C in 1-4% paraformaldehyde, under gentle stirring. Retinas were isolated, stored in methanol at −20°C, and immunostained according to published protocols (29-31). Permeabilized retinas were incubated overnight at 4°C with biotinylated Isolectin B4 (IB4; Vector Laboratories, B-1205, diluted 1:500). Retina were imaged using a Zeiss LSM 780 GaAsp inverted laser scanning fluorescence microscope. Vascular radial extension was quantified by dividing the area labeled by IB4, by the total area of the retina using the Angiogenesis Analyzer plugin on Fiji (http://fiji.sc/Fiji).

### Aortic ring sprouting assay

Aortic ring assay was previously described (32). Briefly, mouse thoracic aortas were isolated and cut into 1 mm-wide rings and embedded in Matrigel® (Corning) and incubated in EBM-2 medium supplemented with VEGF alone or full EGM2 medium containing VEGF, FGF, EGF and IGF-1 (Lonza). Media was replaced every two days. For each sample, the length of eight sprouts at days 6 and 8, originating from the aorta were quantified using the Fiji software (http://fiji.sc/Fiji) from brightfield images taken at 2X.

### Cell culture

Pooled HUVECs (Lonza) were maintained in EGM™-2 (Endothelial Cell Growth Medium-2 BulletKit™;Lonza) at 37°C, 5% CO_2_ and 5% O_2_ as previously described (28). Passages 1 to 8 were used for the experiments described herein.

### Cell migration assay

HUVEC were grown to confluence in a 12-well plate and a scratch wound was created using a sterile p200 pipette tip as previously described (17). Repopulation of the wound in presence of Mitomycin C was recorded by phase-contrast microscopy over 16 hours in a Nikon Ti2-E live-cell microscope. The denuded area was measured at t=0h and t=10h after the wound, and quantified using the ImageJ software (ver. 1.53f51; http://fiji.sc/Fiji). Data were expressed as a ratio of the healed area over the initial wound area.

### Cell proliferation assay

HUVEC were grown at 80% confluence (5×10^3^ cells per well) on glass coverslips in a 24-well plate and starved overnight in serum-free medium (EBM-2, Lonza). They were then incubated for 24 hr in EGM2 containing 10μM BrdU. Immunostaining was performed on cells washed and fixed for 5 min in -20°C methanol, air-dried, rinsed in PBS and permeabilized for 10 min in PBS supplemented with 2% BSA and 0.1% Triton X-100. BrdU positive nuclei were automatically detected in Fijisoftware and normalized to the total number of DAPI-positive nuclei (28).

### Western blotting

Gastrocnemius and soleus muscles were collected and flash-frozen in liquid nitrogen, grinded to power and resuspended in SDS lysis buffer (62.5 mM TRIS pH6,8, 5% SDS, 10 mM EDTA). Protein concentration was determined by DC protein assay (Bio-Rad Laboratories, Reinach, Switzerland), and 10 to 20 μg of protein were loaded per well. Primary cells were washed once with ice-cold PBS and directly lysed with Laemmli buffer as previously described (28, 33). Lysates were resolved by SDS-PAGE and transferred to a PVDF membrane (Immobilon-P, Millipore AG, Switzerland). Immunoblot analyses were performed as previously described (33) using the antibodies described in Table S1. Membranes were revealed by enhanced chemiluminescence (Immobilon, Millipore) using the Azure 280 device (Azure Biosystems) and analyzed using Fiji (ImageJ 1.53c). Protein abundance was normalized to total protein using Pierce™ Reversible Protein Stain Kit for PVDF Membranes (cat 24585; Thermo Fisher Scientific).

### Reverse transcription and quantitative polymerase chain reaction (RT-qPCR)

Pulverized frozen gastrocnemius muscles were homogenized in Tripure Isolation Reagent (Roche, Switzerland), and total RNA was extracted as published (17). After RNA Reverse transcription (Prime Script RT reagent, Takara), cDNA levels were measured by qPCR Fast SYBR™ Green Master Mix (Ref:4385618, Applied Biosystems, ThermoFischer Scientific AG, Switzerland) in a Quant Studio 5 Real-Time PCR System (Applied Biosystems, ThermoFischer Scientific AG, Switzerland), using the primers detailed in Supplementary Table S2.

### Statistical analyses

Data are displayed as means ± standard error of the mean (S.E.M.). Statistical significance was assessed in GraphPad Prism GraphPad Prism 9.1.0 using Student’s t-test, one-way or two-way ANOVA unless otherwise specified. A *p* value of 0.05 or less was deemed statistically significant. Metabolomics data were analyzed using Metaboanalyst software (version 5.0) (34).

## RESULTS

To understand how H_2_S production capacity and the responsible H_2_S producing enzymes are altered as a function of CGL overexpression, we first isolated relevant organs from wild-type (WT) and CGL transgenic mice. We then measured their capacity to produce H_2_S using an enzymatic lead acetate assay (26). As expected, H_2_S production was increased in CGL^Tg^ mice muscle (1.5-fold, Fig. 1A), liver (4 fold, Fig. S1), aorta (Fig. S1) and serum (Fig. S1) samples of as compared to their wild-type counterparts. Next, we measured the mRNA expression of H_2_S-producing enzymes in the gastrocnemius muscle. Consistently, CGL, but not CBS or 3-MST expression was increased (5-fold, Fig. 1B). Immunohistochemical staining of CGL and VE-Cad, an EC marker in gastrocnemius muscle sections confirmed that CGL is mostly expressed by EC and further revealed an increase in CGL levels specifically in EC in the transgenic mice (Fig. 1D).

**Fig. 1:**
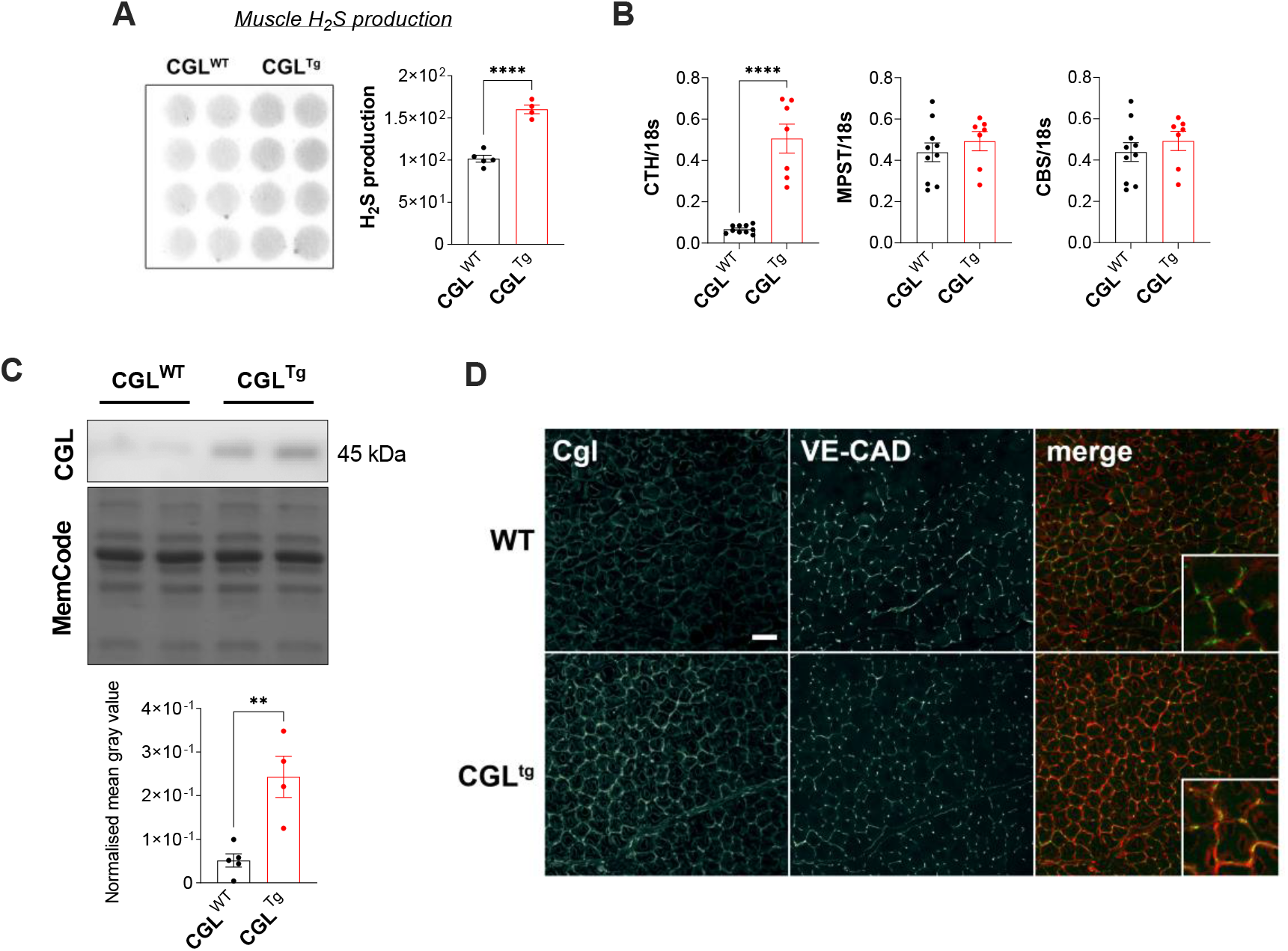
H_2_S production is enhanced in CGL^Tg^ mice gastrocnemius muscle. (A) Measurement of CGL-associated H_2_S production capacity in CGL^WT^ and CGL^Tg^ mice gastrocnemius muscle as detected by lead acetate assay. n= 5 per group (B) Quantitative real-time PCR gene expression analysis of H_2_S-generating enzymes (CTH, MPST and CBS) in gastrocnemius muscle, normalised to 18S expression. n=7-10 per group. (C)CGL protein abundance in gastrocnemius muscle of CGL^WT^ and CGL^Tg^ mice by Western blot. Pierce™ Reversible Protein Stain serves as a loading control. n=4-5 per group. (D) Representative images of gastrocnemius muscle sections (20x) immunostained with CGL and VE-cad antibodies. Data are expressed as mean ± S.E.M. ** p≤0.01 and **** p≤0.0001 by Student’s t-test.

We previously reported that H_2_S/CGL triggers angiogenesis under amino acid restriction (17). Here, we tested whether CGL overexpression alone was sufficient to promote neovascularization in the hindlimb ischemia model, an established *in vivo* model of vascular remodeling via arteriogenesis and angiogenesis (17). Although blood flow was similarly interrupted in both WT and CGL^Tg^ mice immediately after ligation (d0), return of blood flow indicative of neovascularization was accelerated in CGL^Tg^ mice, with significant improvement by d7 after ligation (Fig. 2A). VE-Cad IHC of gastrocnemius muscle sections confirmed a relative increase in capillary density in ischemic legs of CGL^Tg^ mice 21 days post-HLI (Fig 2B). To test the functional significance of CGL overexpression, we performed a treadmill exercise endurance test on d14 after ligation, and observed that the CGL^Tg^ mice ran for a significantly longer time and distance (Fig. 2C). Taken together, our results suggest that CGL-mediated H_2_S production promotes neovascularization and tissue perfusion. Moreover, laminin IHC of gastrocnemius muscle revealed a marked reduction in tissue fibrosis and a concomitant gain in muscle fibre area in CGL^Tg^ mice at day 21 post-HLI (Fig. 2D).

**Fig. 2:**
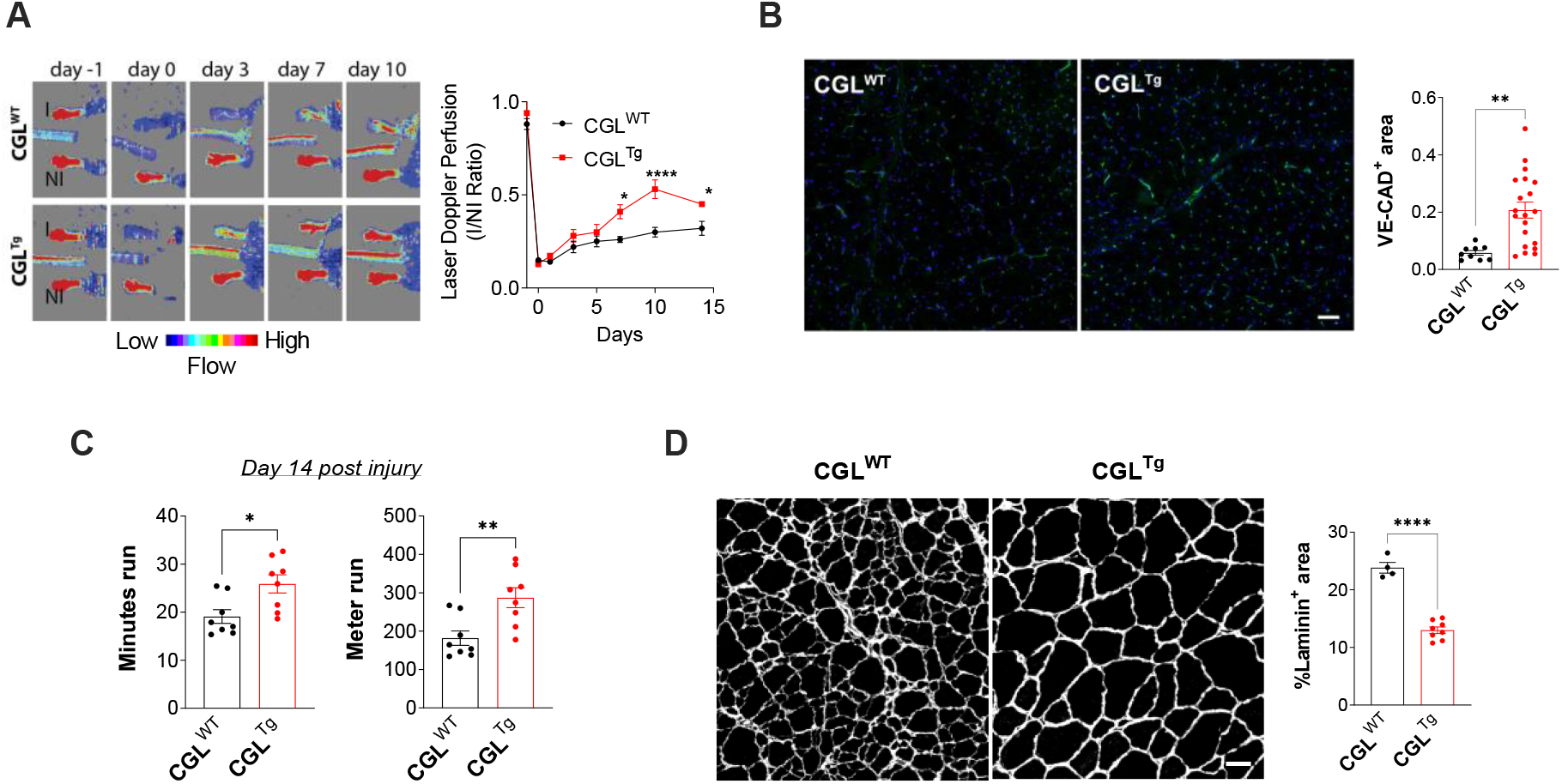
CGL^Tg^ mice shows improved protection from HLI. (A) ZSuperficial perfusion of mouse hindlimb measured by Laser Doppler Perfusion Imaging (LDPI). Results illustrated by representative LDPI images (left) and ratio of LDPI perfusion quantification in ischemic and non-ischemic limbs (right). n=8 per group. (B) Gastrocnemius muscle tissue perfusion 21 days post-HLI as depicted by representative images and quantification (right) of transverse sections stained with VE-cad. n=10-20 per group. Scale bar represent 50μm. (C) Time (left) and distance (right) run on incremental speed test at day 14 post-HLI. n=8 per group. (D) Representative transverse sections (left) and quantification (right) of laminin staining of gastrocnemius muscle at 21 days post-HLI. n=4-8 per group. Scale bar represent 50μm. Data are expressed as mean ± S.E.M. * p≤0.05 and ** p≤0.01 by Student’s t-test.

Next, we sought to better characterise the effects of CGL using a model of sprouting angiogenesis. Vessel sprouting from aortic explants was significantly increased in both VEGF-only and full EGM2 media, with a greater magnitude of increase in VEGF-only media (Fig. 3A, 3B). To assess the effect of CGL overexpression on developmental angiogenesis, we examined the vascular network of whole retina harvested from WT and CGL^Tg^ mice pups at post-natal day 5 (p5). The vascular network appeared identical in both groups, and the radial extension of the network was unchanged (Fig. 3C).

**Fig. 3:**
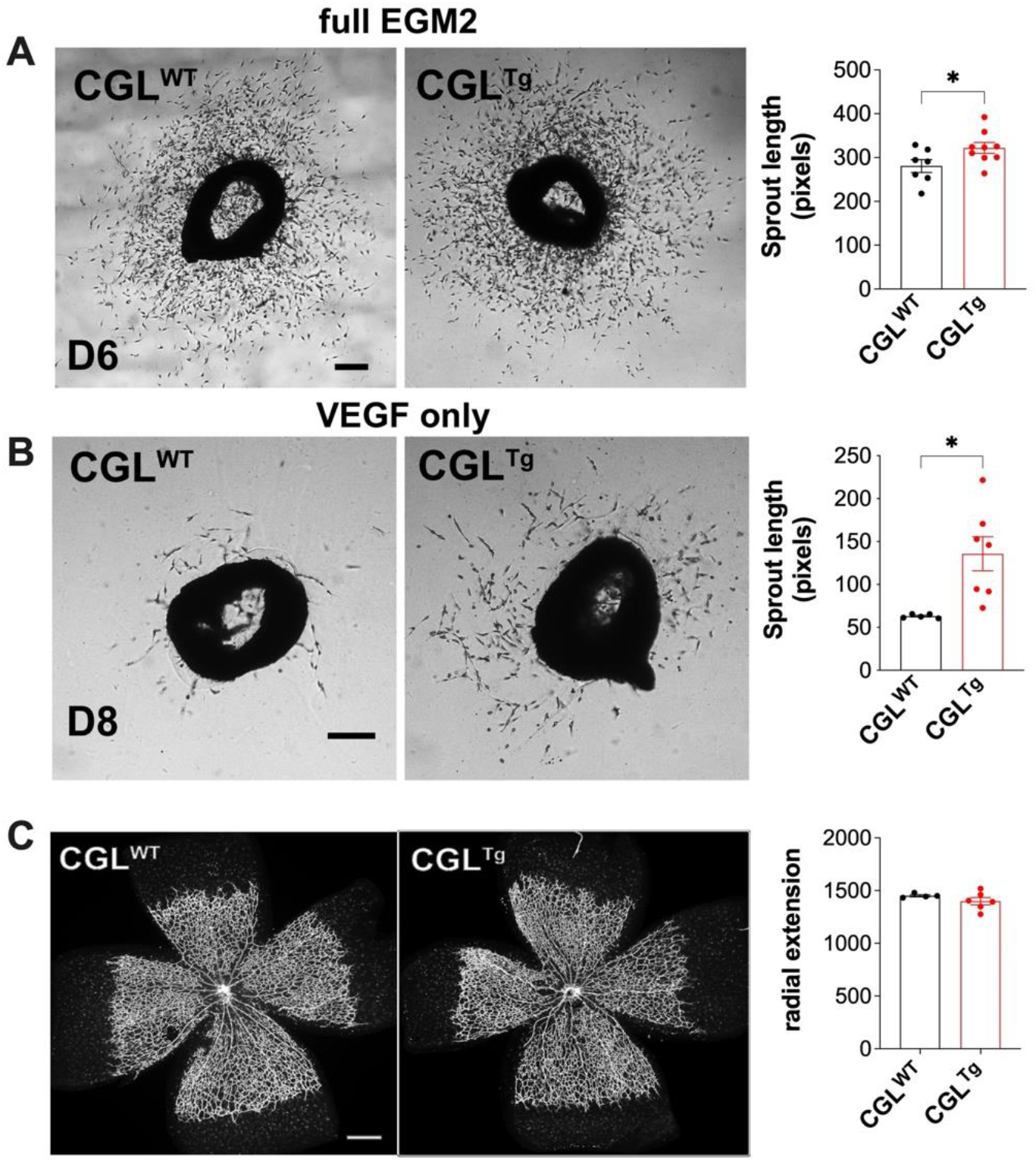
Vascular sprouting is accelerated in CGL^Tg^ mice. (A) Average length of microvessel sprouting from aortic ring explants from CGL^WT^ or CGL^Tg^ mice incubated in full EGM2 media. n=7 per group. Scale bar represents 100μm. (B) Average length of microvessel sprouting from aortic ring explants from CGL^WT^ or CGL^Tg^ mice incubated in VEGF-only EBM2 media. n=7 per group. Scale bar represents 200μm (C) Radial extension of developing mouse retina at post-natal day 5 (p5), with representative images (left) and quantification of radial extension (right) of whole-retina mounts stained with Isolectin B4. n=4-5 per group. Scale bar represent 100μm. Data are expressed as mean ± S.E.M. * p≤0.05 by Student’s t-test.

To elucidate how CGL promoted angiogenesis, we performed targeted metabolomics on baseline (non-ischemic) gastrocnemius from WT and CGL^Tg^ mice. A comparison of global profiles revealed only three significantly differentially abundant metabolites; betaine (0.28 log2 fold change), kynurenic acid (−0.71 log2 fold change) and niacinamide (0.4 log2 fold change), (Fig. 4A and Table S3). The increase in niacinamide was of particular interest due to its role as a key precursor of NAD^+^/NADH.

**Fig. 4:**
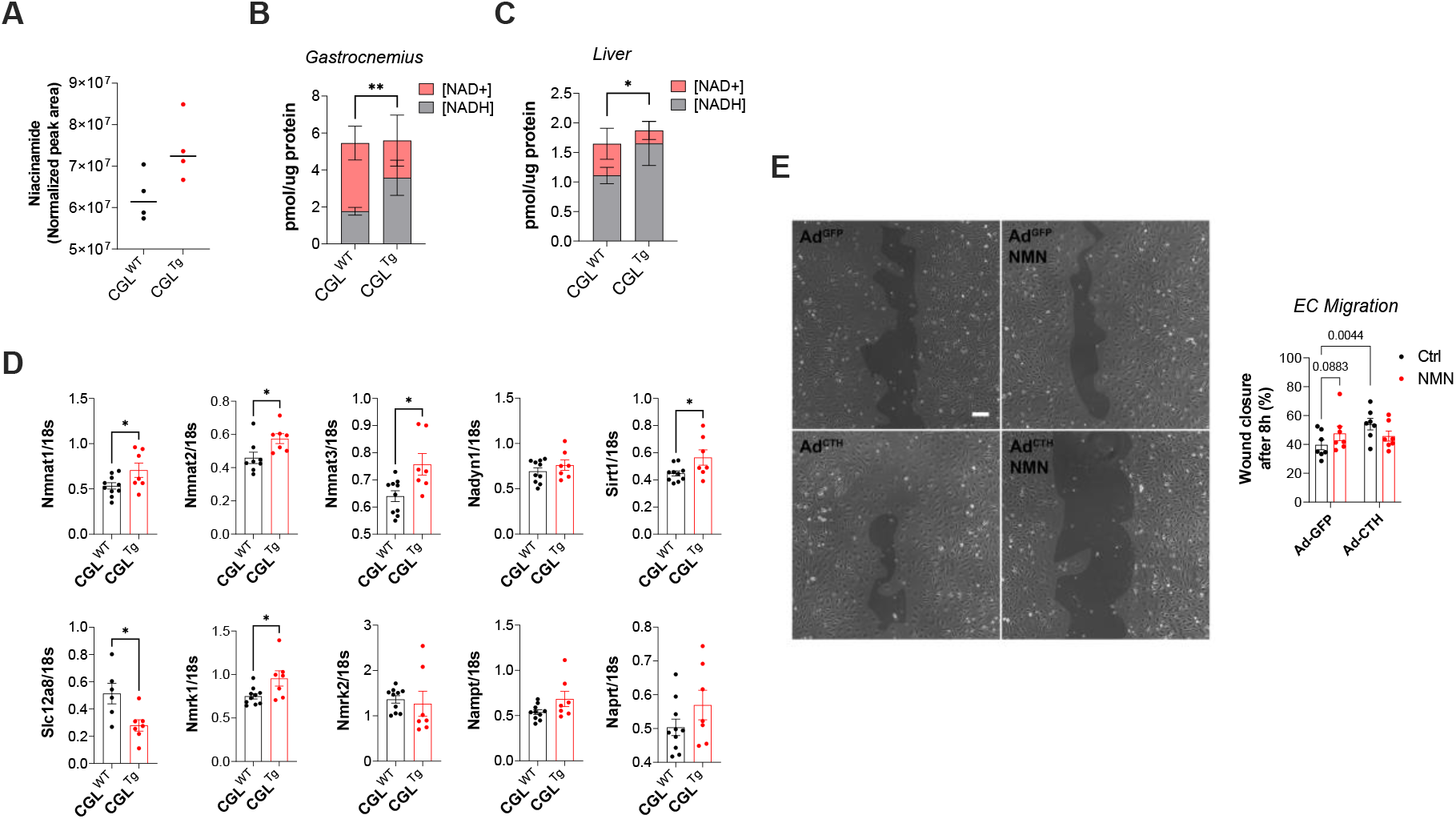
Beneficial effects seen in CGL^Tg^ mice is mediated by NAD^+^/NADH pathways. (A) Niacinamide level in CGL^WT^ and CGL^Tg^ gastrocnemius muscle (n=4). p=0.0569 by Student’s t-test. (B) Quantification of NAD^+^ and NADH concentration (pmol/μg protein) in gastrocnemius muscle (n=5). Data are expressed as mean ± S.D. ** p≤0.01 by Student’s t-test when comparing NADH abundance between groups. (C) Quantification of NAD^+^ and NADH concentration (pmol/μg protein) in liver (n=5). Data are expressed as mean ± S.D. * p≤0.05 by Student’s t-test when comparing NADH abundance between groups. (D) Quantitative real-time PCR gene expression analysis of enzymes associated with the NAD^+^/NADH pathway in gastrocnemius muscle. NAD-generating enzymes: Nmnat1, Nmnat2, Nmnat3, and Nadsyn1. NAD salvage pathway: Slc12a8, Nmrk1, Nmrk2, and Nampt. Preiss-Handler pathway: Naprt. NAD-consuming enzyme: SIRT1. All genes were normalised to 18S expression. n=7-10 per group. (E) Endothelial cell migration across scratch with representative image (left, 10x magnification) and quantification (right) of HUVECs infected with GFP-or CGL-expressing adenovirus, ± NMN supplementation as indicated. n=7 per group. Scale bar in representative images represent a length of 100 μm. n=7 per group. Data are expressed as mean ± S.E.M. * p≤0.05 by Student’s t-test.

Of importance, H_2_S dependent increases in capillary density during exercise was previously shown to involve NAD^+^ (20). NADH levels were increased two-fold in the gastrocnemius muscle of CGL^Tg^ mice, while NAD^+^ was equally decreased. (Fig. 4B). These findings were recapitulated in the liver (Fig. 4C). We performed qPCR analyses on a series of genes closely associated with the NAD^+^/NADH system. It revealed significant increases in Nmnat1, Nmnat2 and Nmnat3 RNA abundance; significant genes in the NAD^+^ salvage pathway (Fig. 4D). Moreover, we see an increased expression of SIRT1 (Fig. 4D), an NAD^+^ consuming enzyme important in various cellular processes.

To understand the effects of boosting the NAD^+^/NADH system upon aspects of angiogenesis, we transfected HUVECs with GFP-or CGL-expressing adenovirus, in presence or absence of nicotinamide mononucleotide (NMN), a NAD^+^ precursor. Both CGL overexpression and NMN treatment individually increased HUVEC migration. However, the combination of the two did not result in further improvement, suggesting a redundant effect on migration (Fig 4E). CGL overexpression or NMN treatment had no effect on cell proliferation (Fig. S2).

## DISCUSSION

Here, we show a beneficial role of CGL overexpression in a murine model of PAD. In addition, CGL overexpression modulates the NAD^+^/NADH biosynthesis pathway, which improves EC migration and thus angiogenesis and recovery from hindlimb ischemia.

In this study, we observed an accelerated improvement in limb perfusion and capillary density following ischemia in mice with CGL overexpression, while also seeing a marked improvement in functional recovery measured by endurance running capability. This builds on our previous findings whereby the increase of CGL-associated H_2_S production boosts VEGF-dependent angiogenesis (17). Our *in vivo* findings build on previous work (11, 13, 35) suggesting that H_2_S is beneficial for neovascularization of the ischemic hindlimb. Interestingly, we observed a fascinating interplay between H_2_S and the NAD^+^/NADH system. Thus, the NAD^+^/NADH precursor niacinamide was increased in the muscle of CGL-overexpressing mice, and the NAD^+^/NADH balance shifted. Moreover, the expression of several key enzymes within the NAD^+^/NADH biosynthesis pathway changed, suggesting a robust response to changes in H_2_S production. Prior *in vitro* studies saw a transient increase in NAD^+^ concentration in HUVECs exposed to the H_2_S donor NaHS (20). Here, NAD^+^/NADH ratio was decreased in the gastrocnemius of CGL-overexpressing mice. Our measurements of steady state metabolite levels limit our interpretations, as there could be differences in rate of NAD turnover, or a difference in the shorter-term versus longer-term effects of H_2_S augmentation. Further experiments using flux measurements with stable isotope labelled substrate will be required to determine this definitively.

A possible explanation for this observation is related to the observation that the H_2_S-mediated improvement in ischemic limb neovascularization stems from an enhancement of glycolysis (17, 36). Glycolysis involves a series of enzymatic reactions catalysed by GAPDH and LDH and uses NAD^+^ as a coenzyme (19). Thus, the increase in relative abundance of NADH in the CGL^Tg^ muscle may represent an increase in glycolysis-related NAD^+^ consumption. We previously demonstrated that CGL overexpression in EC increase glycolysis and ATP production (17). Thus, it is likely that NADH accumulation is driven by NAD+ consumption, which is known to play a key role in neovascularization in the hindlimb ischemia model (20). Despite our novel observations, our mechanistic understanding of the interplay between H_2_S and the NAD^+^/NADH system is still incomplete and warrants further investigation.

Importantly, CGL-overexpressing mice muscle feature increased SIRT1 gene expression. Endothelial cell-specific SIRT1 knockout mice have impaired neovascularization after hindlimb ischemia (37), and SIRT1 has been shown to help EC integrate pro-angiogenic signals secreted by ischemic myocytes (20). Thus, the increased SIRT1 expression seen in CGL overexpressing mice likely supports neovascularization. As a NAD^+^-consuming enzyme, SIRT1 increased expression could contribute to the reduced NAD^+^ concentration in CGL overexpressing mouse muscle.

On a cellular level, we found that boosting H_2_S and NAD^+^/NADH production enhances EC migration but not proliferation *in vitro*. This is consistent with prior studies which highlighted substantial improvements in endothelial cell migration with H_2_S supplementation (17, 38). In addition, boosting both H_2_S and NAD^+^/NADH pathways simultaneously did not have a synergistic effect on EC migration, suggesting that the pro-migratory effects of H_2_S and NAD^+^/NADH occur through the same mechanism. The enhancement of EC migration was previously shown to be required for the endothelial tip cells of the sprouting vasculature (39). Improvements in endothelial tip cell migration would lead to faster development of vascular structures – ultimately leading to improved neovascularization of the ischemic area.

Although our study mainly explored the changes within the NAD^+^/NADH pathways with CGL overexpression, we would like to highlight that our metabolomics analysis revealed two other significantly differentially abundant metabolites, namely, betaine (0.28 log2 fold change) and kynurenic acid (−0.71 log2 fold change). Betaine is part of the transulfuration pathway, allowing regeneration of methionine from homocysteine via the betaine homocysteine methyltransferase (9). Betaine accumulation is consistent with a shift toward utilization of homocysteine to generate cysteine via CGL. Accumulation of kynurenic acid suggest a swift in the kynurenine pathway, which outlines the conversion of tryptophan to NAD^+^. Of note, most enzymes in this pathway are dependent on vitamin B6 pyridoxal 5’-phosphate (PLP), similarly to the transulfuration pathway enzymes CGL and CBS. Accumulation of kynurenic acid could be due to reduced kynurenine pathway activity due to NAD+ accumulation of increased PLP utilization by CGL. Interestingly, both the betaine and tryptophan/kynurenic acid pathways are important in the regulation of immune cells. Previous studies has shown that macrophages are polarised towards the pro-repair M2-phenotype in presence of betaine (40). Moreover, the same phenomena was observed in the context of indoleamine 2,3-dioxygenase overexpression, a key enzyme driving the catabolism of tryptophan to kynurenic acid (41). The M2 macrophage phenotype have been shown to be vital in the neovascularisation in the context of the hindlimb ischemia model (42), and could hence posit a plausible mechanism of action to explain the improvements in neovascularisation when CGL is overexpressed.

Although our findings have given us cause to be optimistic, we are aware of its limitations. We endeavoured to replicate PAD pathology with our hindlimb ischemia procedure. Although it may be the gold standard in replicating the pathology *in vivo*, it does not reflect the chronic nature of PAD development. Of note, we recently demonstrated that H_2_S production capacity and plasma sulfide concentrations were reduced in patients with PAD (15). The chronicity of PAD may allow for the development of compensatory adaptations, not be present in our hindlimb ischemia model. Furthermore, we have yet to fully characterize potential off-target effects of CGL overexpression. We could anticipate changes in the relative levels of specific amino acid pools, in particular sulfur-containing amino acids, and it is unclear what implications this may have on ischemic adaptations. Although we see increases in NAD precursors and in many NAD^+^-related genes, our steady state metabolomic measurements limit our ability to determine rates of NAD^+^ flux or fate of NAD^+^. Future studies using stable isotope tracing are required to test our hypotheses. Our study only draws conclusions upon the effects of the global increase in CGL, but it remains to be seen whether the protection is due to enhanced CGL production locally within EC at the ischemic site, or whether it is due to distal systemic CGL overexpression. As our long-term goal is to offer therapeutic angiogenesis, it would therefore be important to identify the cell types and organ systems that drive the pro-angiogenic phenotype as targets for the production of H_2_S.

---

**Fig. S1:**
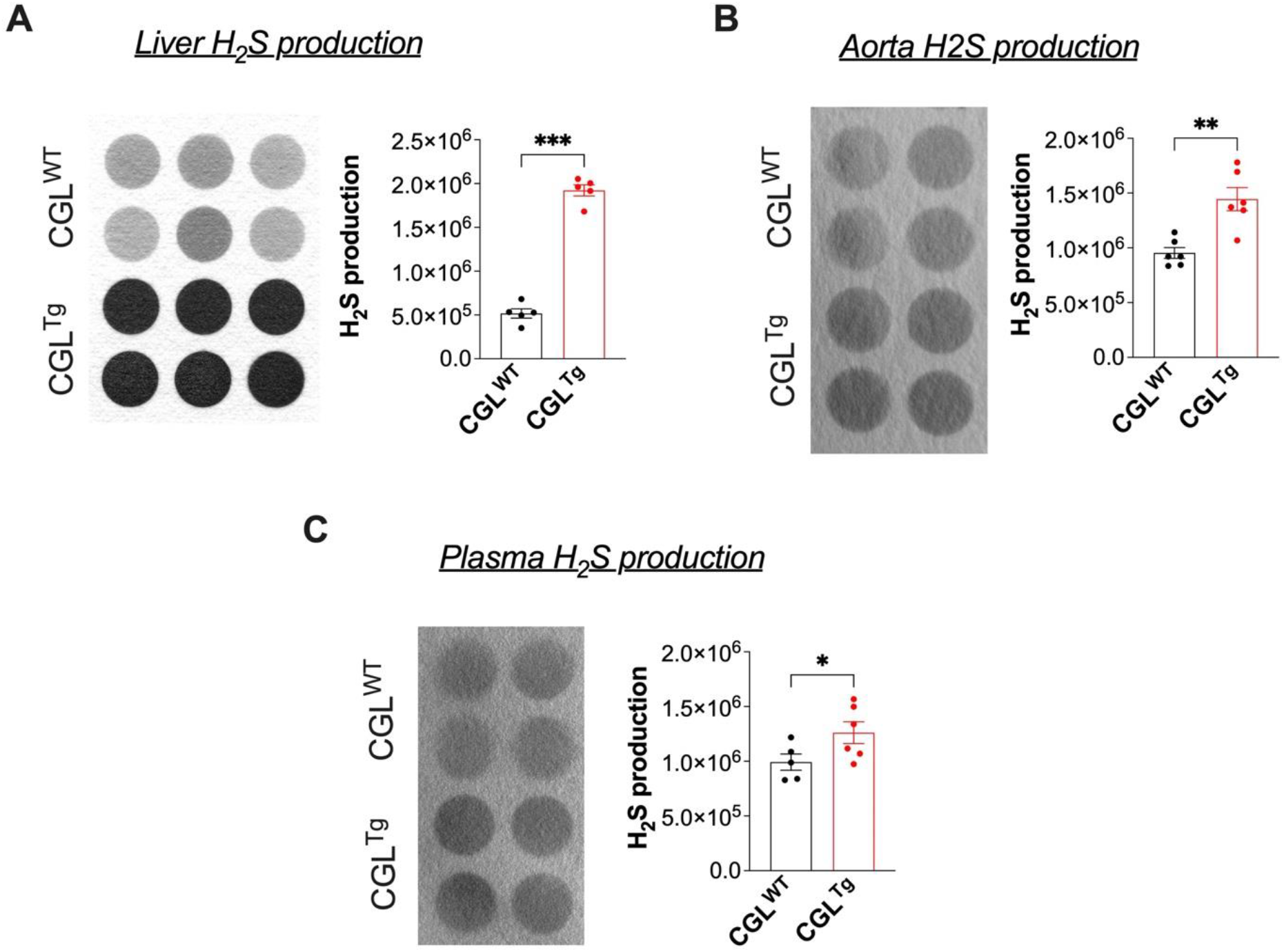
CGL^Tg^ mice produce greater amounts of H_2_S in various organs. (A) CGL-associated H_2_S production capacity in CGL^WT^ and CGL^Tg^ mice liver as detected by lead acetate assay. n=5 per group. (B) CGL-associated H_2_S production capacity in CGL^WT^ and CGL^Tg^ mice aorta as detected by lead acetate assay. n=6 per group. (C) CGL-associated H_2_S production capacity in CGL^WT^ and CGL^Tg^ mice plasma as detected by lead acetate assay. n=6-7 per group. Data are expressed as mean ± S.E.M. * p≤0.05, **p<.01;*** p≤0.001 by Student’s t-test.

**Fig. S2:**
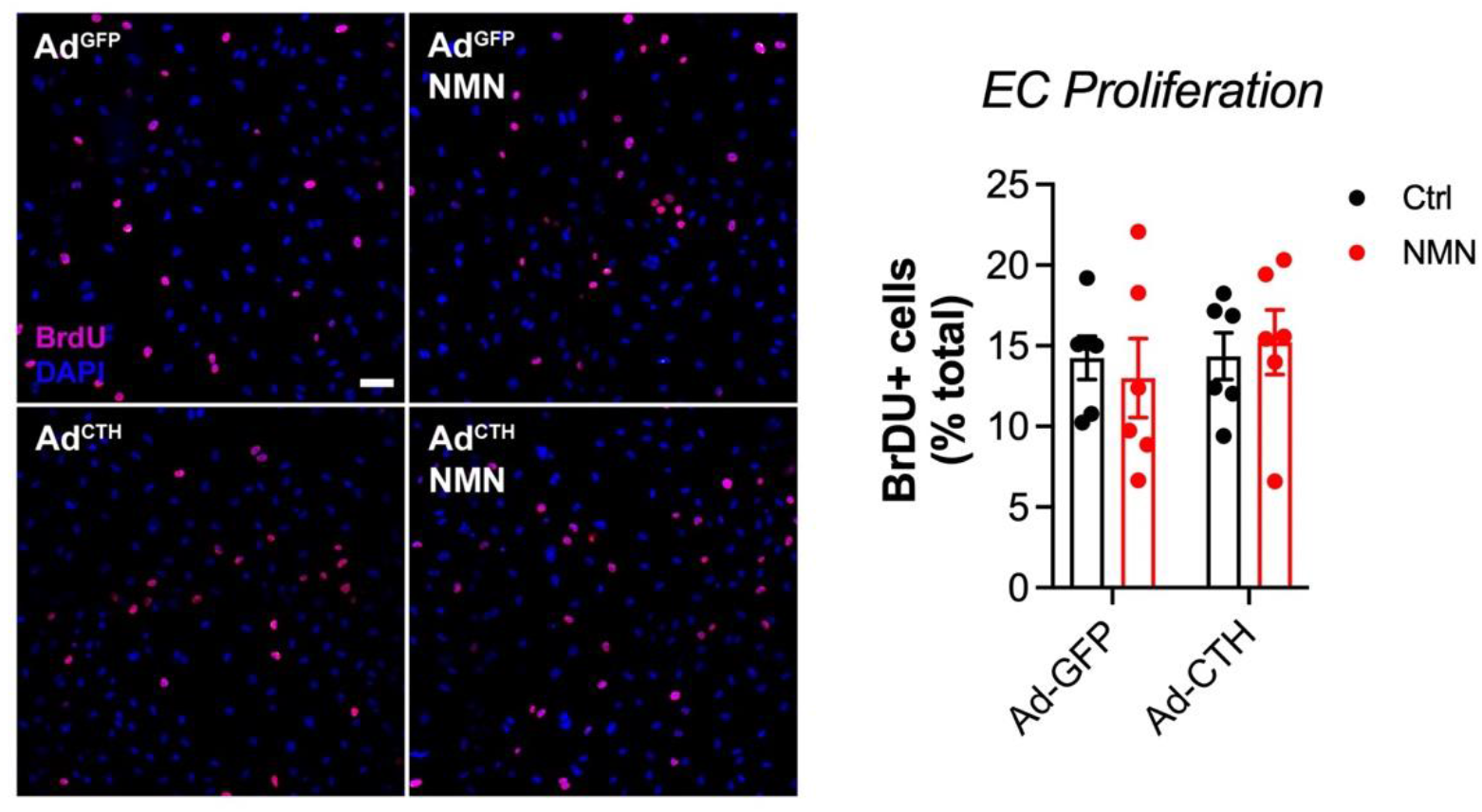
EC proliferation capacity unchanged when NAD or H_2_S pathways are enhanced. (A) Proliferation capability of HUVECs transfected with GFP-or CGL-expressing adenovirus, ± NMN supplementation as indicated. Scale bar in representative images represent a length of 50 μm. n=6 per group. Data are expressed as mean ± S.E.M. no statistical differences as assessed by Student’s t-test.

**Supplementary Table S1:** Antibodies.

| **Target antigen** | **Vendor** | **Catalog #** | **Working concentration** |
| --- | --- | --- | --- |
| Laminin | Sigma | L9393 | 1/200 |
| Ve-Cadherin | Abcam | AB33168 | 1/100 |
| CGL | ProteinTech | 12217-1-AP | 1/1000 (WB)<br>1/100 (IHC) |
| Anti-Rabbit HRPO | Thermo Fisher Scientific | 31460A21109 | 1/20000 (WB)<br>1/250 (ICC) |
| Anti-mouse HRPO | Jackson ImmunoResearch Labs | 115-035-146 | 1/15000 (WB) |
| Anti-Rabbit HRPO | Thermo Fisher Scientific | 31460 | 1/20000 (WB) |
| BrdU | BD Biosciences | 555627 | 1/200 (ICC) |
| Goat anti-Rabbit IgG Secondary Antibody, Alexa Fluor 680 | Thermo Fisher Scientific | A21109 | 1/500 |
| Goat anti-Rabbit IgG Secondary Antibody, Alexa Fluor 405 | Thermo Fisher Scientific | A31556 | 1/500 |
| Goat anti-Rat IgG Secondary Antibody, Alexa Fluor 488 | Thermo Fisher Scientific | A11006 | 1/500 |
| Donkey anti-Rabbit IgG Secondary Antibody, Alexa Fluor 488 | Thermo Fisher Scientific | A21206 | 1/500 |

**Supplementary Table S2:**
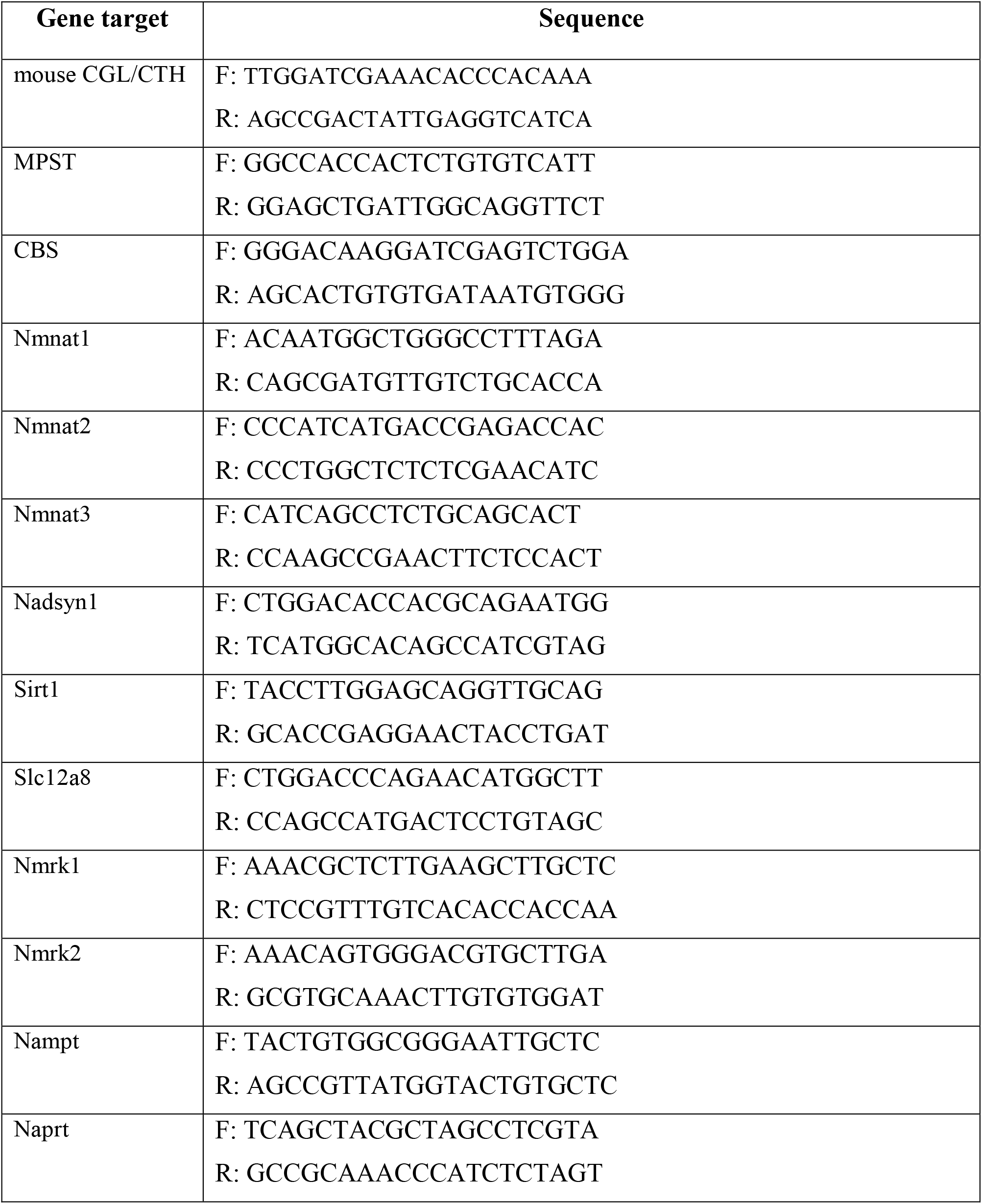
DNA oligo primers.

